## Supplemental Figures for "Comparative analysis of the syncytiotrophoblast in placenta tissue and trophoblast organoids using snRNA sequencing"

### Supplementary Figures

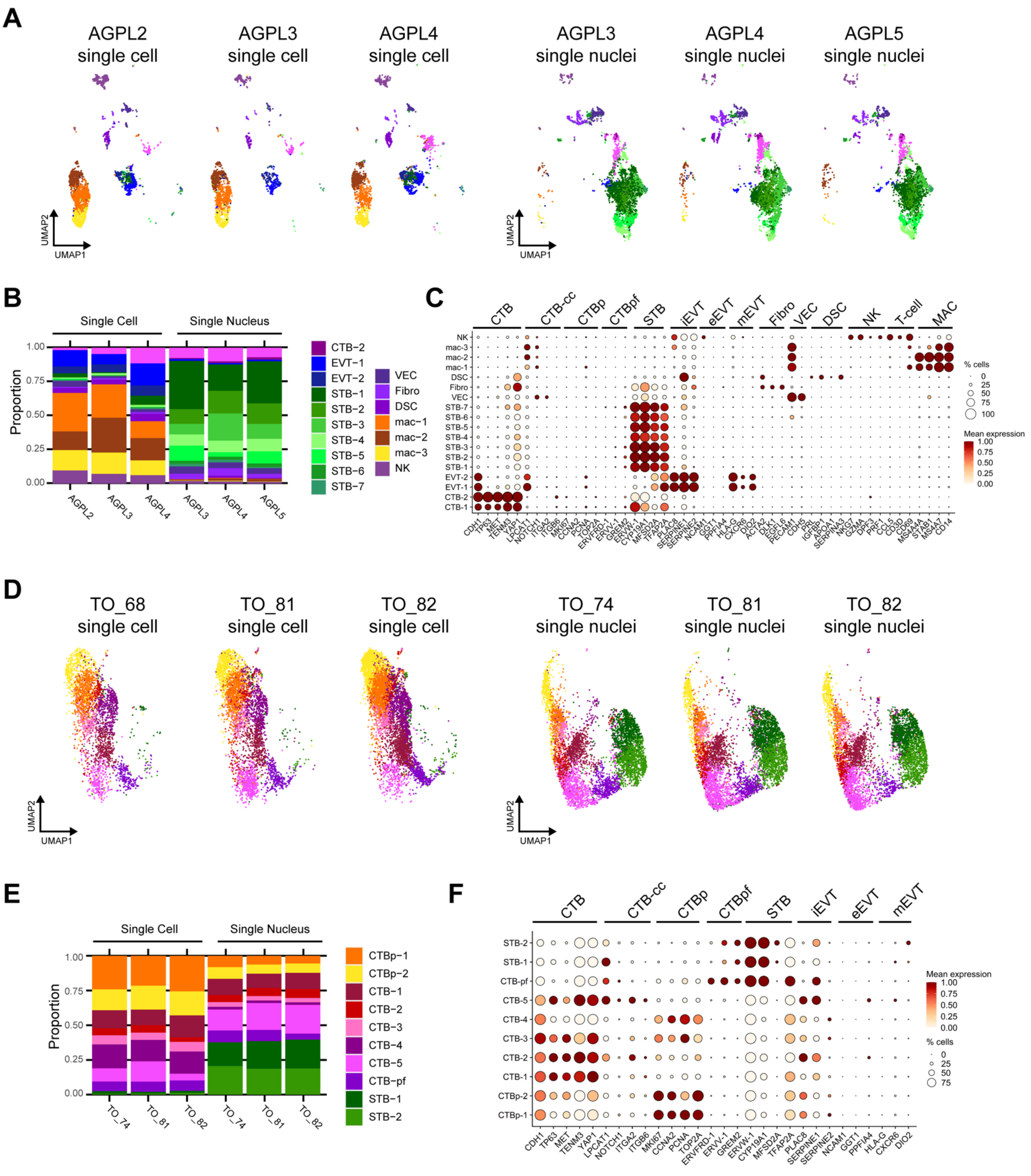


**Supplemental Figure 1.1**: **Characterization of cell/nucleus types in the integrated SC/SN dataset.** Each patient (3 tissues) and sequencing approach (SC or SN) for the integrated SC/SN full-term tissue dataset was individually plotted on a UMAP in **A**, cell/nucleus type proportions are plotted as a barplot in **B**, and dot plot of gene expression for the marker genes of each cell/nucleus type shown in **C**. Each TO line and sequencing approach (SC or SN) for the integrated SC/SN TO dataset was individually plotted on a UMAP in **D**, cell/nucleus type proportions are plotted as a barplot in **E**, and dot plot of gene expression for the marker genes of each cell/nucleus type shown in **F**. For the dot plots in **C** and **F**, the size of the dot demonstrates the percent of cells/nuclei expressing a given gene and color represents the mean expression value. The labels on the top of the graph refer to the established marker genes used to identify the identity of each cluster. Cell/nucleus types are annotated as follows: cytotrophoblasts (CTB), cytotrophoblast pre-fusion (CTB-pf), syncytiotrophoblast (STB), extravillous trophoblast (EVT), dendritic stem cell (DSC), vascular endothelial cell (VEC), fibroblast (Fib), natural killer cell (NK), and macrophage (Mac). Cell/nucleus types that contained multiple clusters are identified with -number after the cell name.

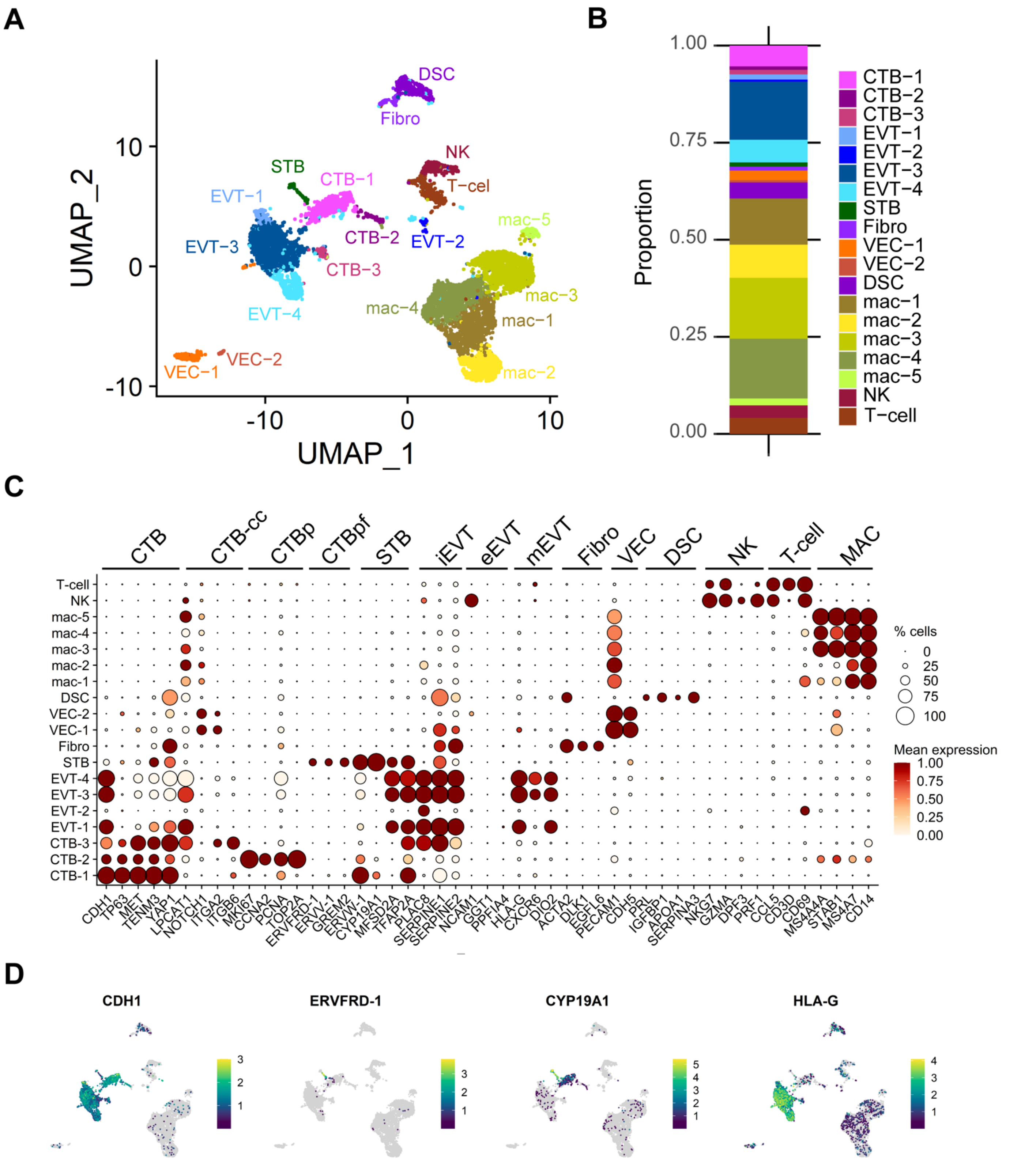


**Supplemental Figure 1.2: Characterization of cell types in the full-term tissue SC dataset. A** UMAP visualization of cell types found via Louvain clustering in Seurat. **B** Barplot demonstrating proportions of each cell type in the dataset. **C** Dot plot visualization of the gene expression of each cell type. The size of the dot demonstrates the percent of cells expressing a given gene and color represents the mean expression value. The labels on the top of the graph refer to the established marker genes used to identify each cluster. **D**

Featureplots of gene expression of specific marker genes for CTB (CDH1), CTB-pf (ERVFRD-1), STB (CYP19A1), and HLA-G (EVT). Cell types are annotated as follows: cytotrophoblasts (CTB), cytotrophoblast pre-fusion (CTB-pf), syncytiotrophoblast (STB), extravillous trophoblast (EVT), dendritic stem cell (DSC), vascular endothelial cell (VEC), fibroblast (Fib), natural killer cell (NK), and macrophage (Mac). Cell types that contained multiple clusters are identified with -number after the cell name.

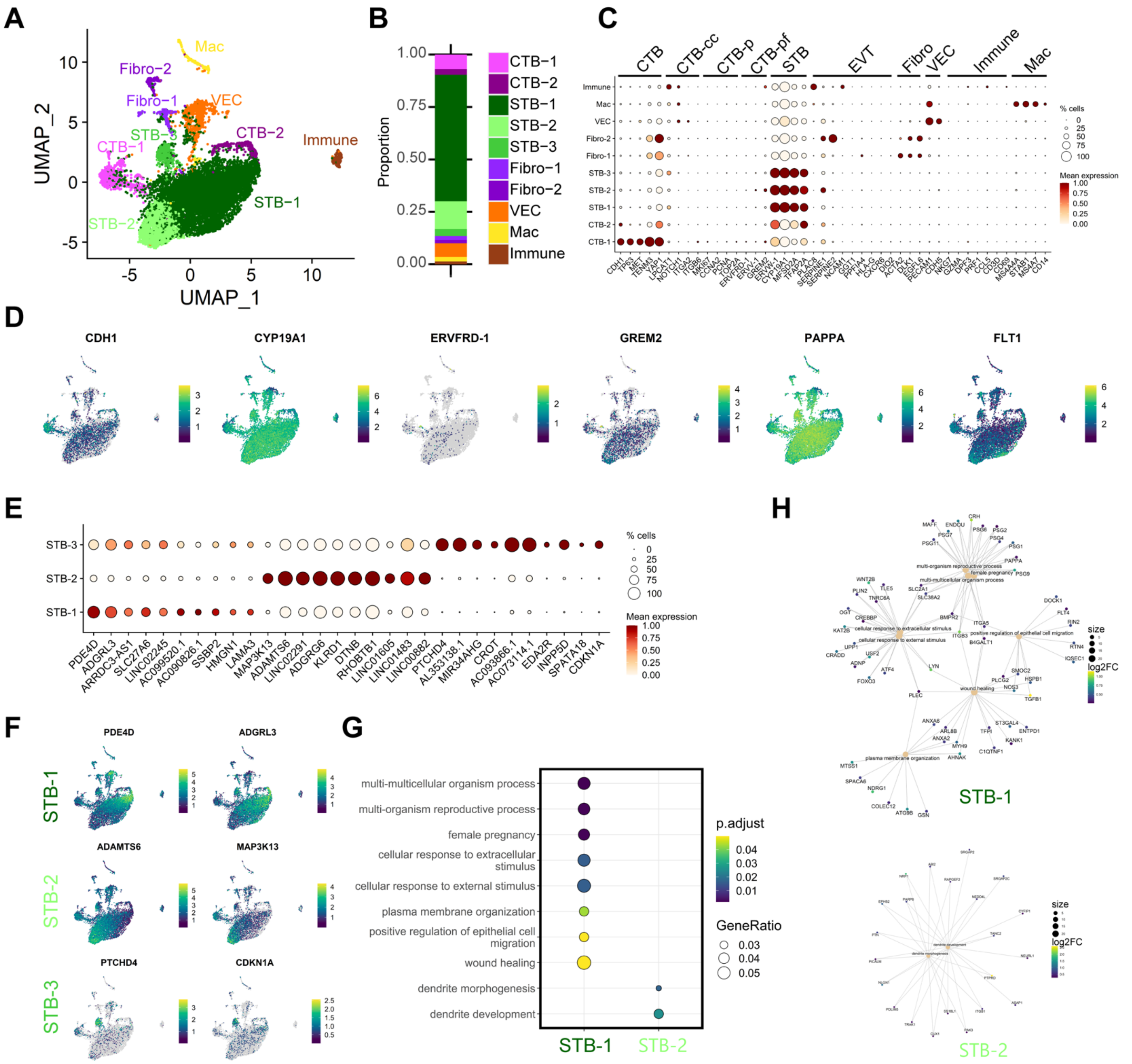


**Supplemental Figure 1.3: Characterization of nucleus types in tissue SN dataset. A** UMAP visualization of nucleus types found via Louvain clustering in Seurat. **B** Barplot demonstrating proportions of each nucleus type in the dataset. Nucleus types are annotated as follows: cytotrophoblasts (CTB), cytotrophoblast pre-fusion (CTB-pf), syncytiotrophoblast (STB), vascular endothelial cell (VEC), fibroblast (Fibo), macrophage (Mac), unidentified immune cell (Immune). Nucleus types that contained multiple clusters are identified with -number after the cell name. **C** Dot plot visualization of the gene expression of each nucleus type. The size of the dot demonstrates the percent of nuclei expressing a given gene and color represents the mean expression value. The labels on the top of the graph refer to the established marker genes used to identify the identity of each cluster. **D** Featureplots of gene expression of specific marker genes for CTB (CDH1), STB (CYP19A1), CTB-pf

(ERVFRD-1 and GREM2). In addition, markers for the two terminally differentiated nucleus subtypes found in Wang et al., were visualized (PAPPA and FLT1). **E**. Top ten differentially expressed genes (DEGs) of each STB subtype is shown as a dotplot. **F** Featureplots of selected genes in E enriched in each STB-subtype. **G** Biological GO terms associated with the DEGs of each subtype were found and plotted with ClusterProfiler. No GO terms were statistically associated with STB-3. **H** The genes associated with each GO term in G are plotted as a CNET plot in ClusterProfiler. The size of the GO term node is scaled by the number of associated genes (scale bar to right) while each gene is colored with the relative log2FC gene expression.

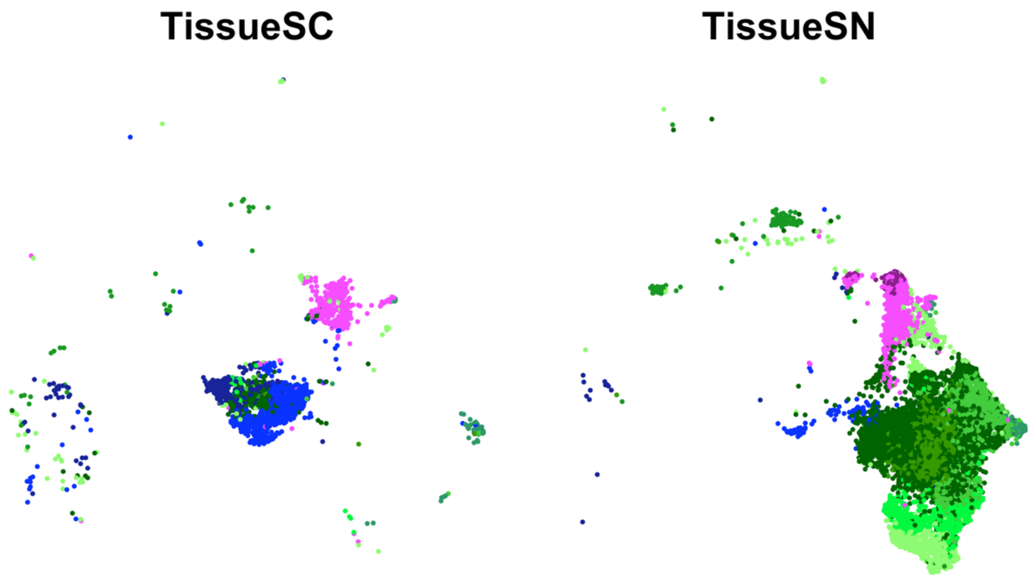


**Supplemental Figure 1.4** UMAP of the integrated SC and SN dataset separated into individual UMAPs by single cell or nucleus processing type for primary tissue. Only the trophoblast cell types are included with EVTs in blue, CTBs in pink, and STBs in green.

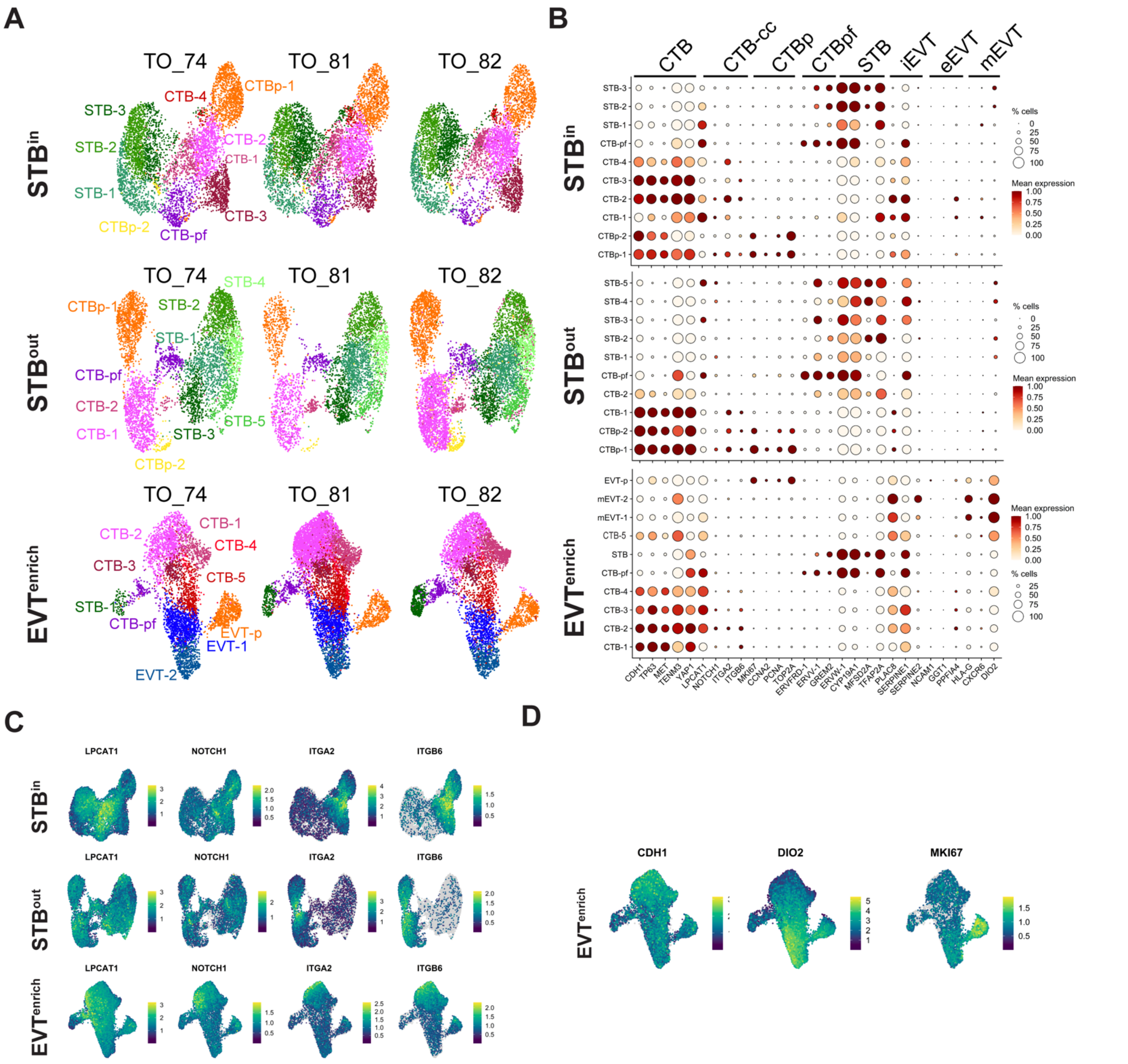


**Supplemental Figure 2.1: Characterization of nucleus subtypes in individual TO datasets. A** Each TO cell line was individually plotted on a UMAP for the integrated SN STB$^{in}$, STB$^{out}$, or EVT$^{enrich}$ datasets. **B** Dot plot of gene expression for the marker genes of each nucleus type. The size of the dot demonstrates the percent of nuclei expressing a given gene and color represents the mean expression value. The labels on the top of the graph refer to the established marker genes used to identify the identity of each cluster. **C** Feature Plot demonstrating gene expression of the CTB-CC markers in each TO culture condition. **D** Feature Plot demonstrating gene expression of a CTB marker (CDH1), an EVT marker (DIO2), and a mitotic marker (MKI67).

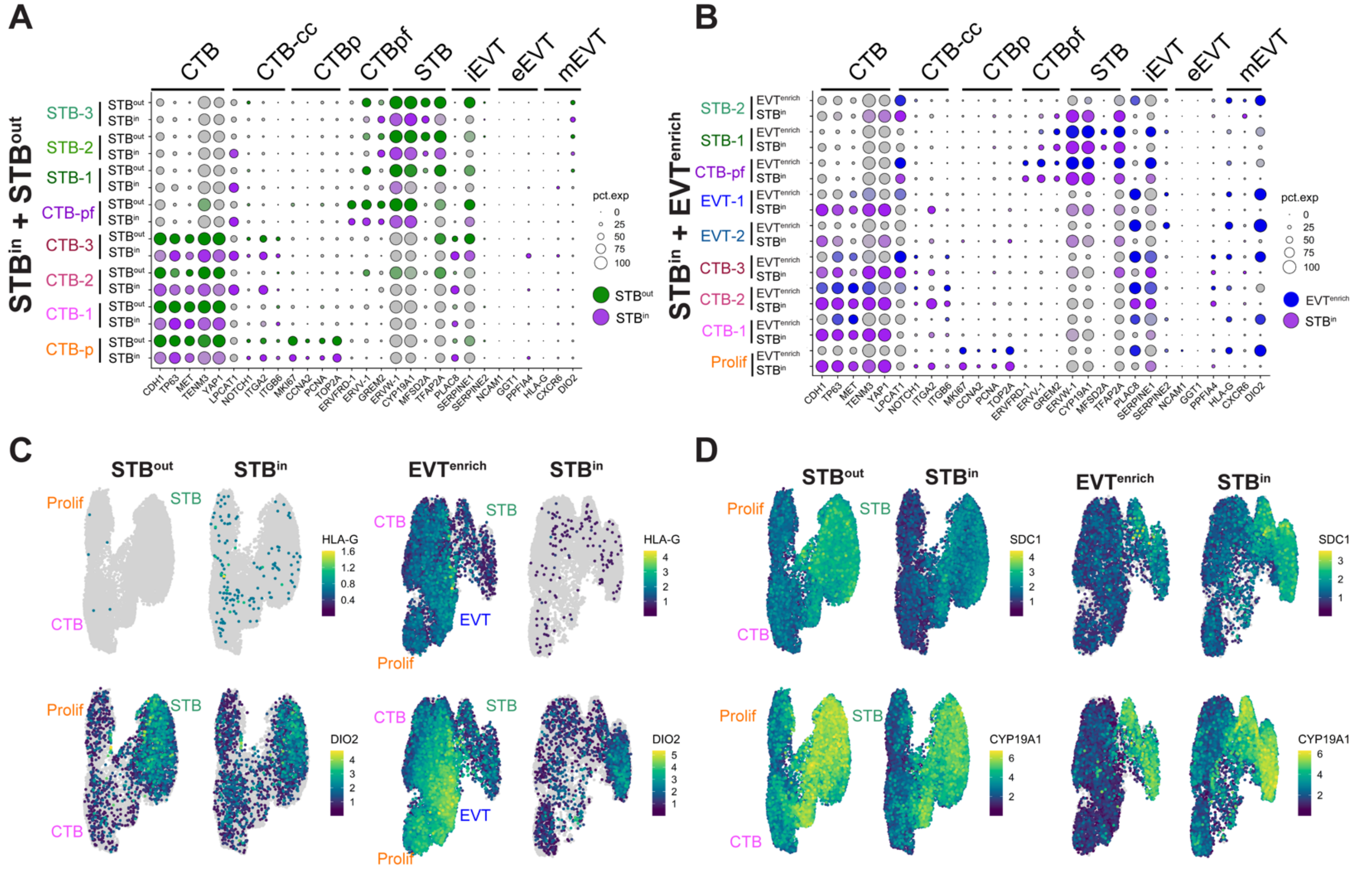


**Supplemental Figure 2.2: Characterization of nucleus types in the merged TO SN datasets.** Dot plot of gene expression for the marker genes of each nucleus type for the integrated STB$^{in}$ + STB$^{out}$ dataset (**A**) or STB$^{in}$ + EVT$^{enrich}$ dataset (**B**). Individual gene expression is shown for each nucleus type of each culture condition and labels are found on the left-hand axis of the graph. The size of the dot demonstrates the percent of nuclei expressing a given gene and color represents the mean expression value. The labels on the top of the graph refer to the established marker genes used to identify the identity each cluster. Feature Plot of integrated datasets split by culture condition of EVT markers (HLA-G and DIO2, **C**) or STB markers (SDC1 and CYP19A1, **D**).

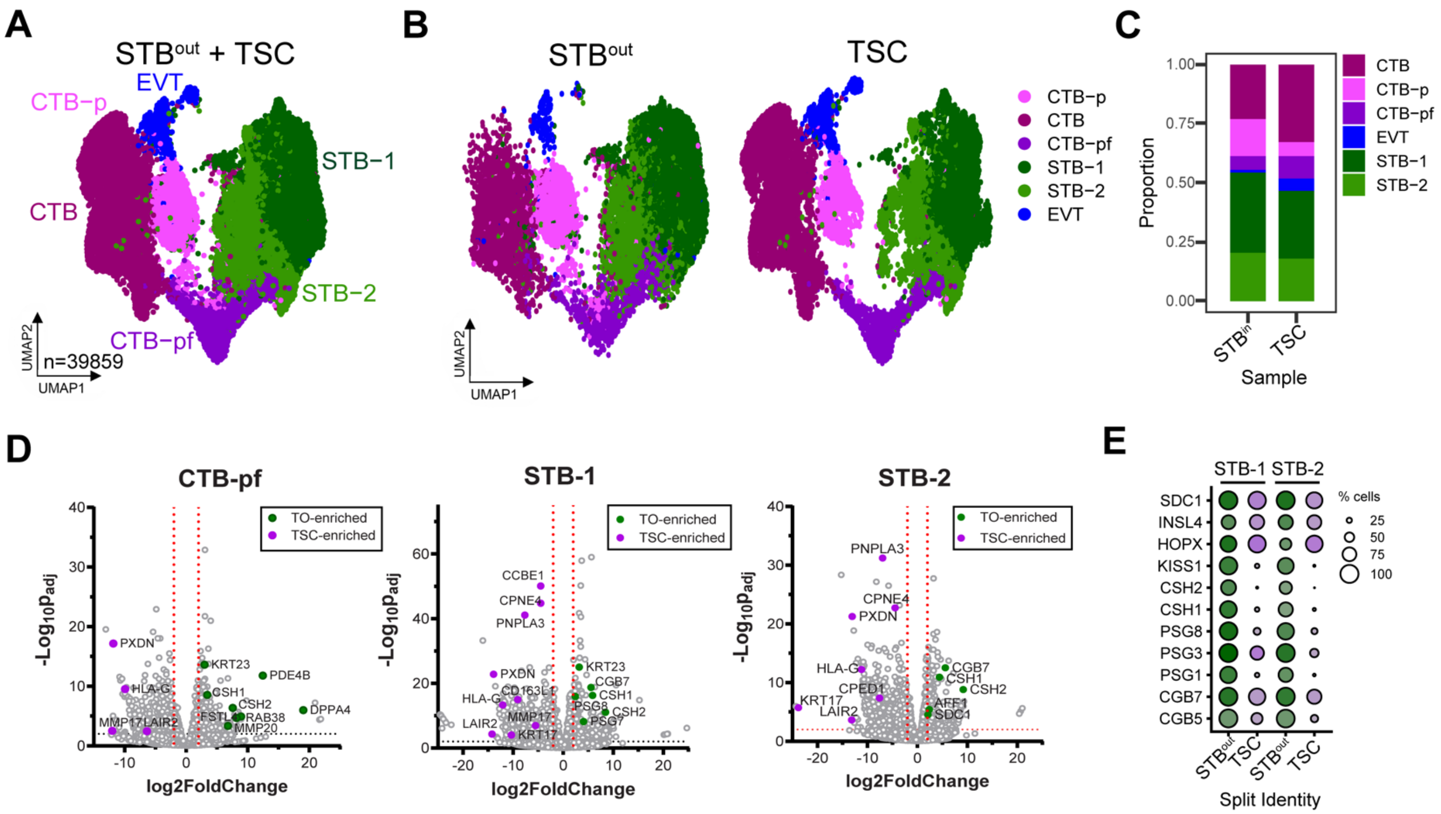


**Supplemental Figure 2.3: Comparison of TO and TSC SN datasets.** UMAP of integrated STB$^{out}$ and TSC dataset (**A**) or plotted individually (**B**) with nucleus types found via Louvain clustering in Seurat and pseudocolored. **C** Barplot demonstrating proportions of each nucleus type in the integrated dataset. **D** DEseq2 was used to find differentially expressed genes between TO and TSC datasets in CTB-pf, STB-1, and STB-2 clusters and plotted as a volcano plot. Select representative genes are highlighted for TO (green) or TSC (purple). **E** Dot plot of individual genes in the STB-1 and STB-2 subtypes split into either TSC or TO populations.

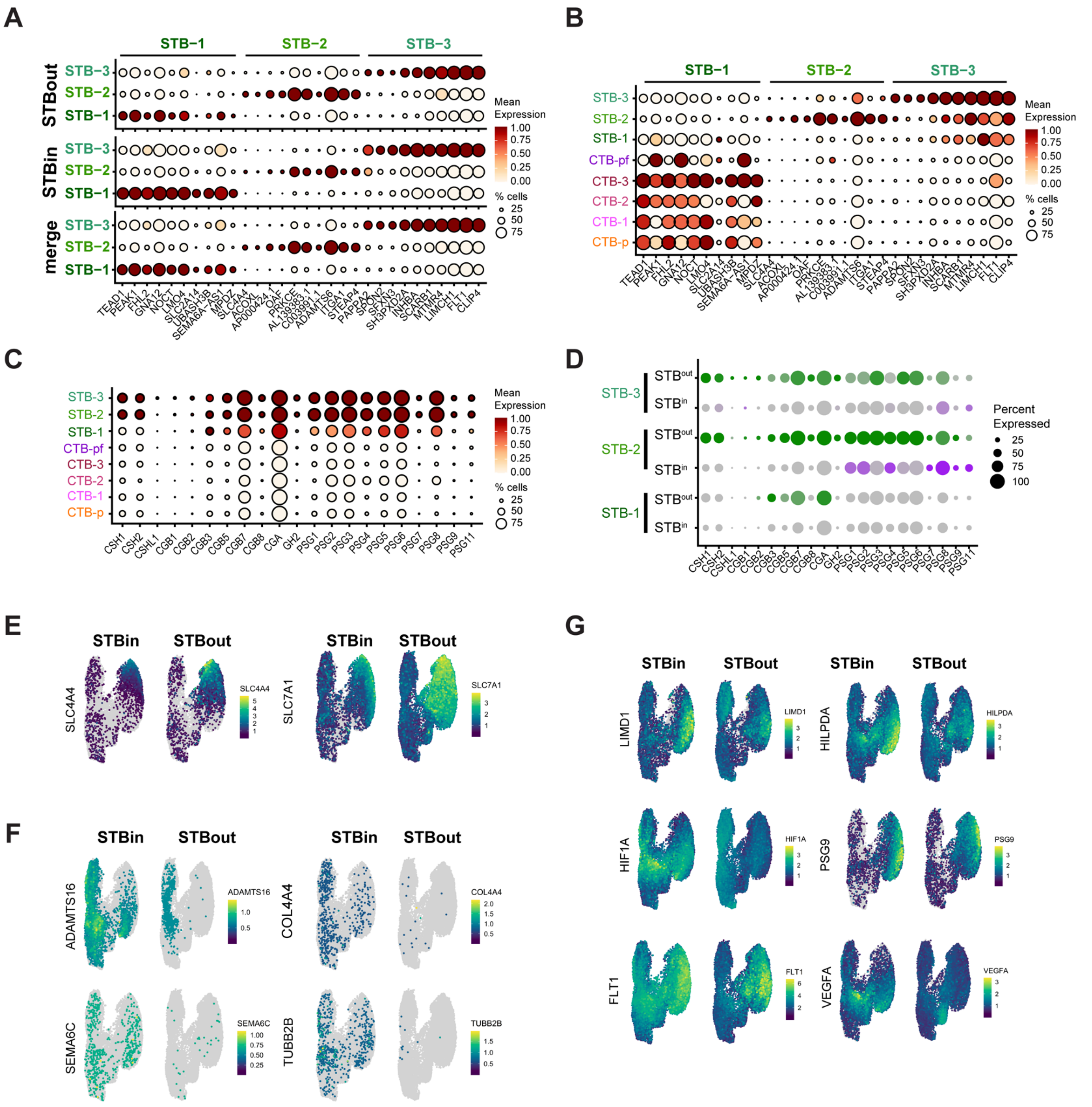

**Supplemental Figure 3.1: STB subtype analysis and expression differences in STB$^{in}$ vs STB$^{out}$ TOs. A** Top ten differentially expressed genes (DEGs) of each STB subtype was determined in the integrated STB$^{in}$+STB$^{out}$ dataset comparing just the three STB clusters and plotted for either STB$^{out}$, STB$^{in}$, or the merged dataset. The labels on the top of the graph refer to the top DEGs from each STB subtype. **B** The same genes found in A were plotted for every cluster in the STB$^{in}$+STB$^{out}$ dataset to demonstrate the expression of STB-1 marker genes in the CTB clusters. **C** Dotplot demonstrating the mean expression of key STB pregnancy hormones in each nucleus type of the integrated STB$^{in}$+STB$^{out}$ dataset. **D** The same hormones plotted in A were plotted as a dot plot and split into either $STB^{out}$ (green) or $STB^{in}$ (purple) expression. For each dotplot in A-D The size of the dot demonstrates the percent of nuclei expressing a given gene and color represents the mean expression value. **E** Featureplot demonstrating expression of transporter proteins enriched in STB-3. **F** Feature plot demonstrating expression of extracellular matrix genes enriched in $STB^{in}$. **G** Feature plot of the hypoxia associated genes LIMD1, HIF1A, and FLT1.

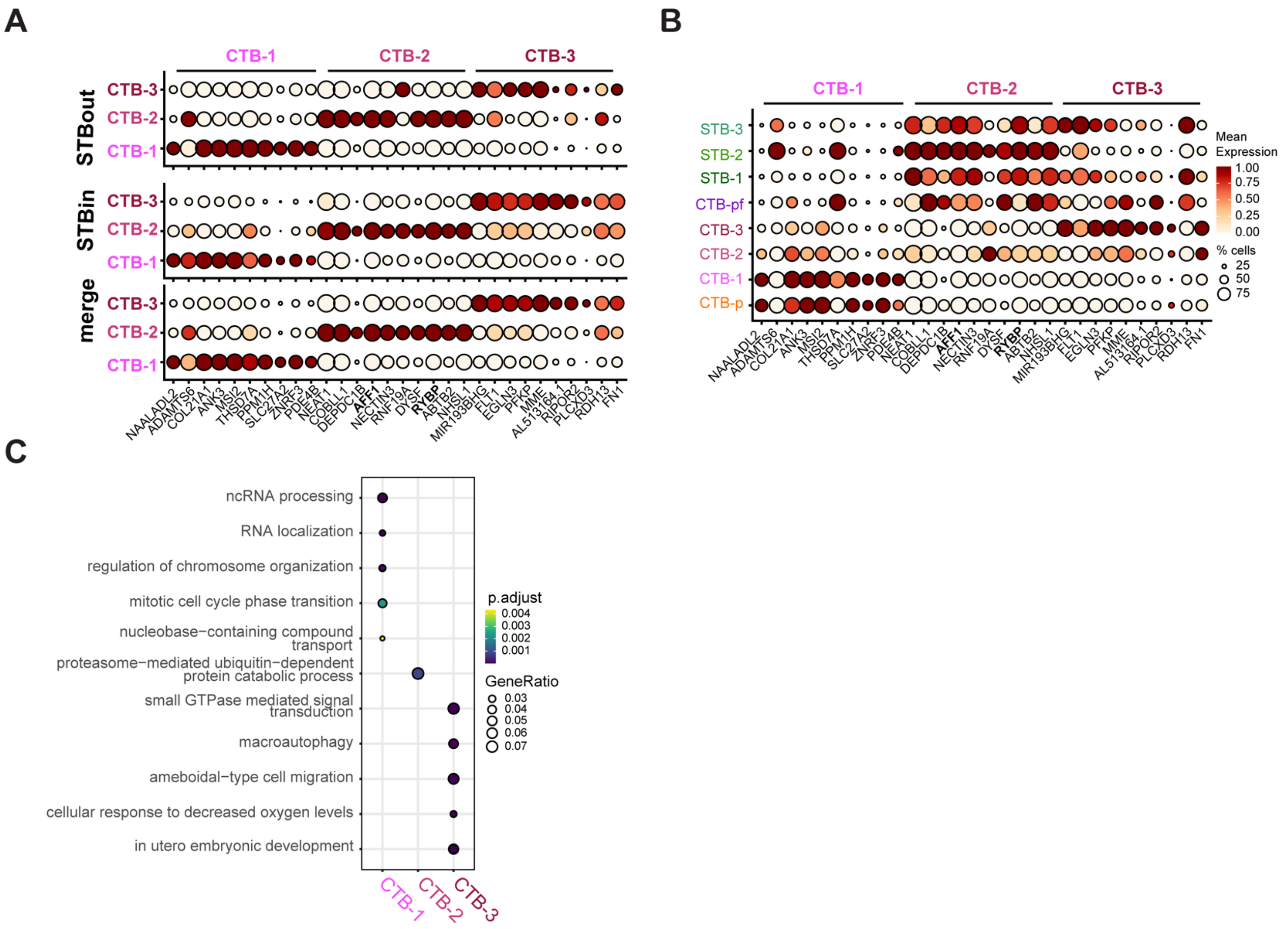


**Supplemental Figure 3.2: CTB subtype analysis and expression differences in STB$^{in}$ vs STB$^{out}$ TOs. A** Top ten differentially expressed genes (DEGs) of each CTB subtype was determined in the integrated STB$^{in}$+STB$^{out}$ dataset comparing just the three CTB clusters and plotted for either STB$^{out}$, STB$^{in}$, or the merged dataset. The labels on the top of the graph refer to the top DEGs from each CTB subtype. **B** The same genes found in A were plotted for every cluster in the STB$^{in}$+STB$^{out}$ dataset to demonstrate the expression of CTB-2 marker genes in the STB clusters. **C** Top Biological Process GO terms for each CTB subtype determined with clusterProfiler.

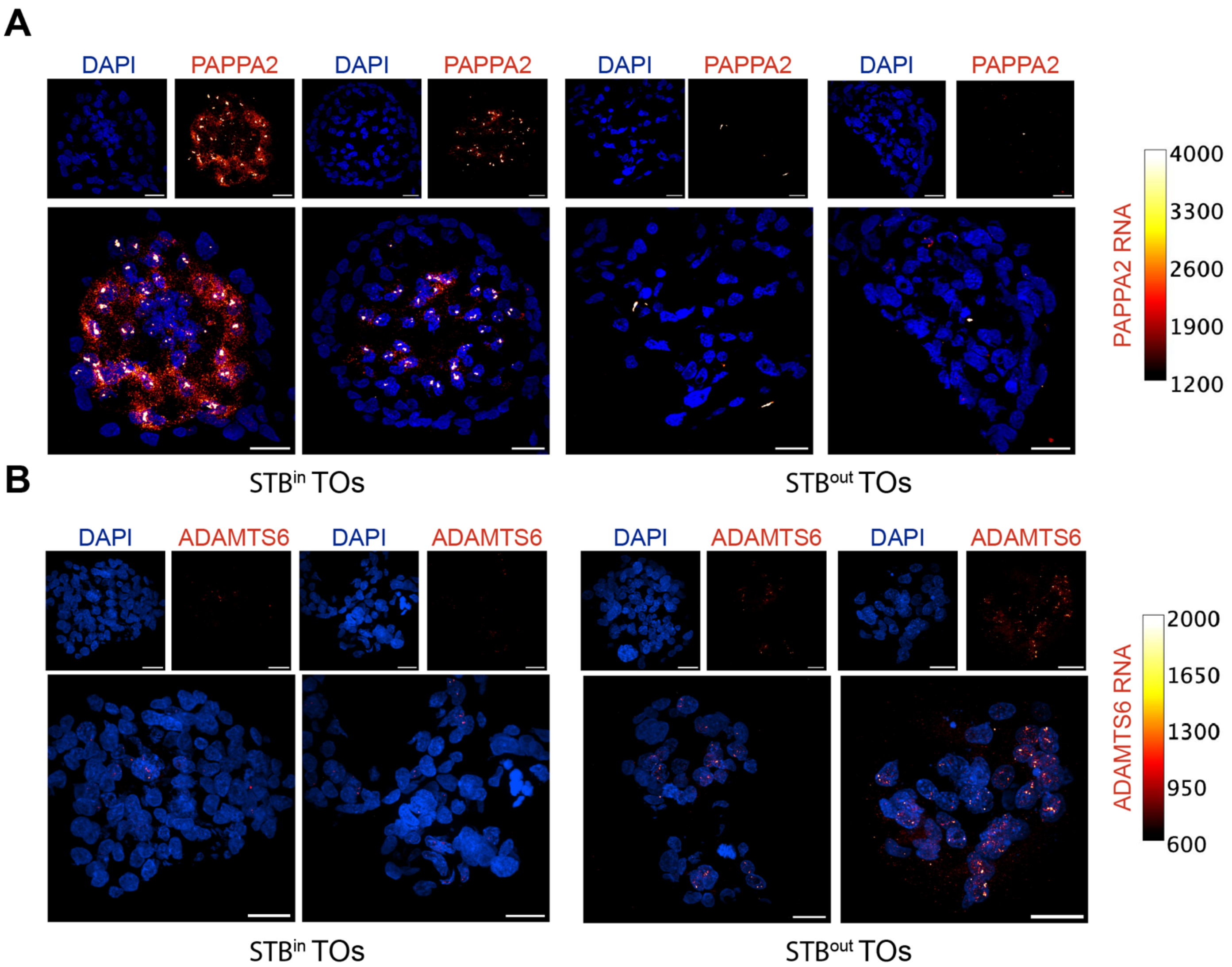


**Supplemental Figure 3.3: Representative images of RNA FISH in TOs.** RNA fluorescence in situ hybridization (FISH) was performed on STB$^{in}$ and STB$^{out}$ TOs against PAPPA2 in **A** and ADAMTS6 in **B**. Cells were stained with DAPI and visualized in blue while RNA expression is represented as the "Red Hot" color in FIJI and the calibration color bar is shown to the right of each image. Images were identically thresholded for RNA expression, but DAPI thresholding was adjusted independently in each image to account for differences in nuclear brightness between STB$^{in}$ and STB$^{out}$ TOs. Each scale bar represents 20μm.

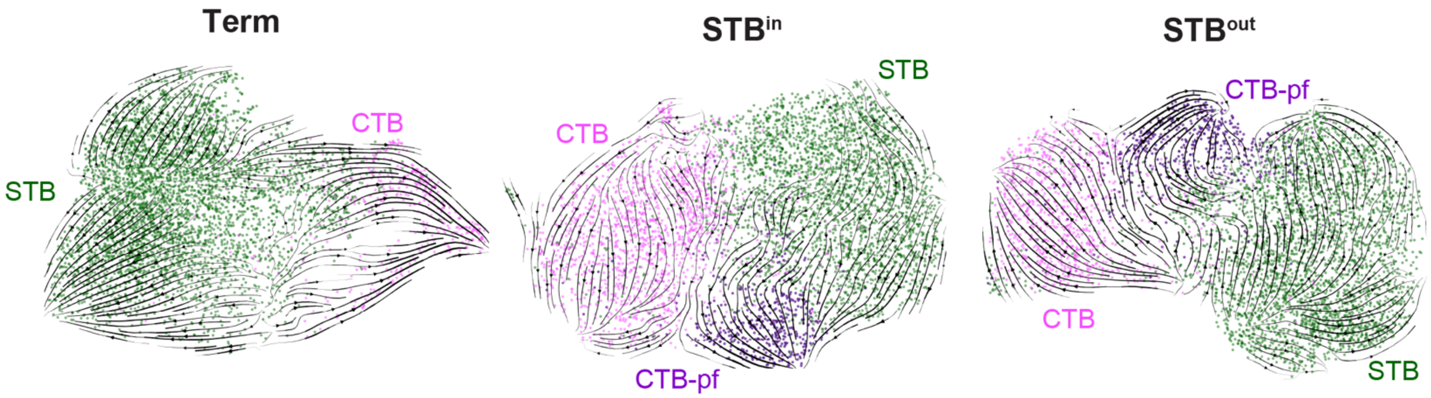


**Supplemental Figure 4.1: RNA velocity traces.** Velocity derived from scVelo is visualized as streamlines on UMAP for either full-term tissue (Term), STB$^{in}$, or STB$^{out}$ datasets.

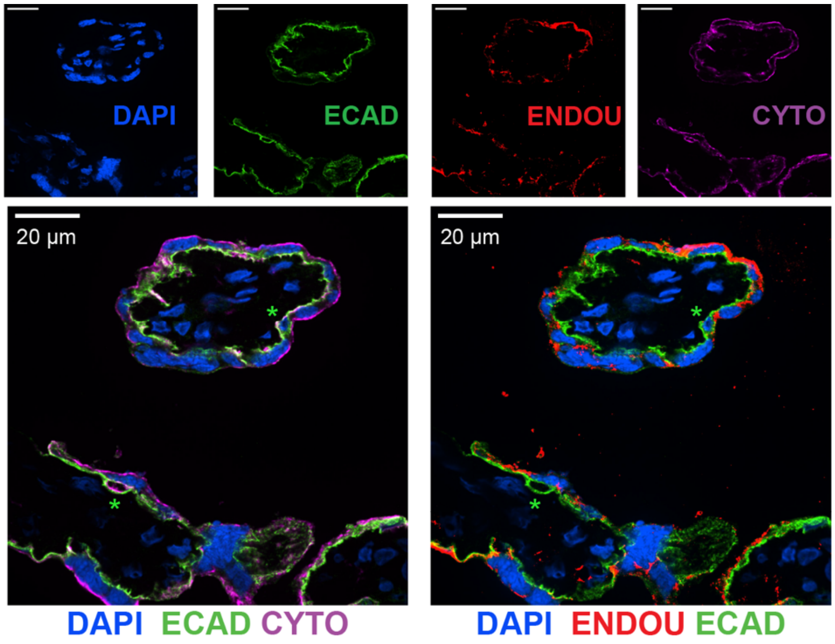


**Supplemental Figure 5.1 The STB marker ENDOU demonstrates E-cadherin and cytokeratin-7 can designate CTB and STB nuclei in tissue.** Immunofluorescence of full-term tissue was performed with E-cadherin (ECAD), cytokeratin (CYTO), and ENDOU and stained with DAPI to label nuclei. Green stars represent CTB cells.

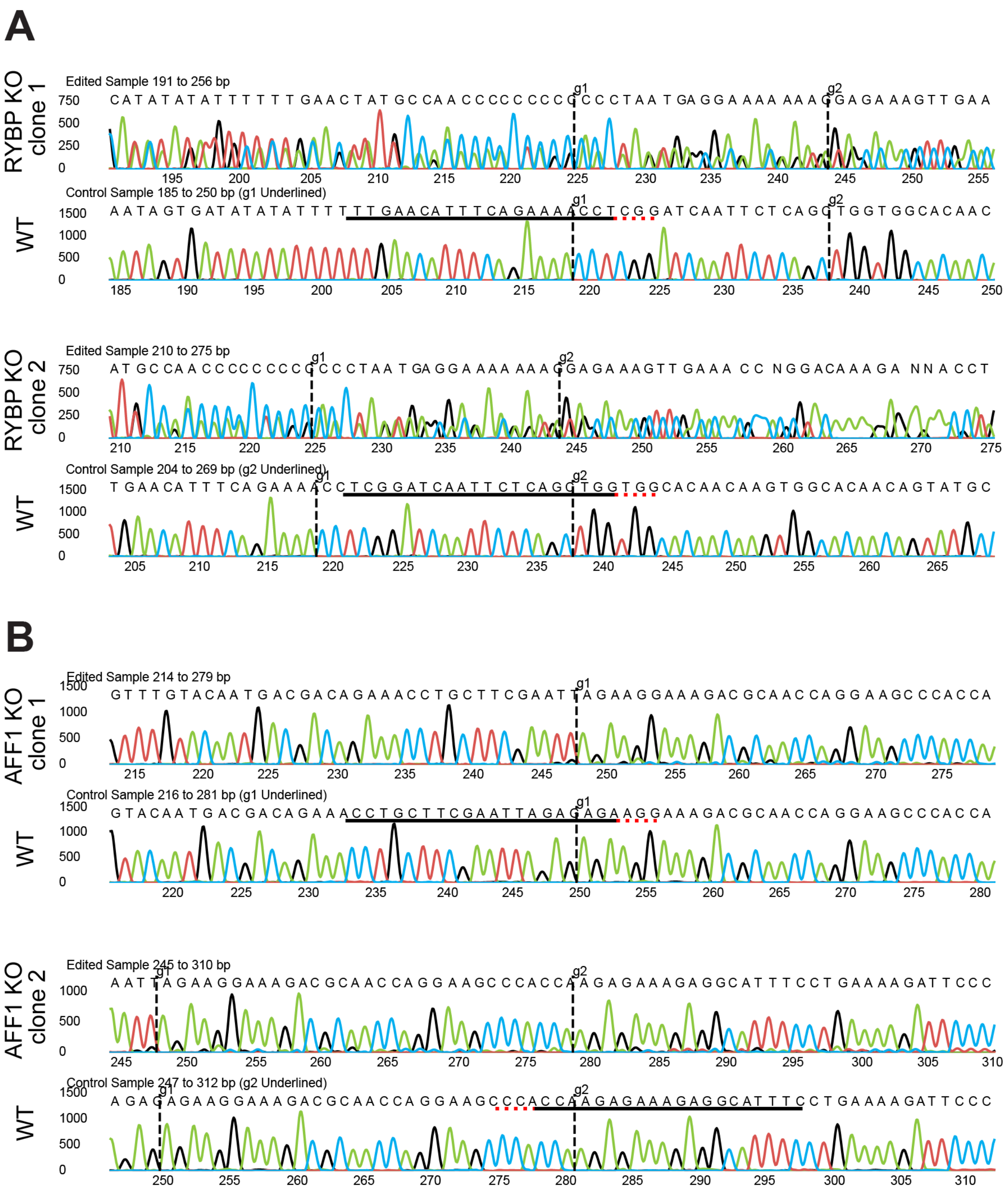


**Supplemental Figure 5.2 Sanger sequencing validation of RYBP and AFF1 KO TO clones.**

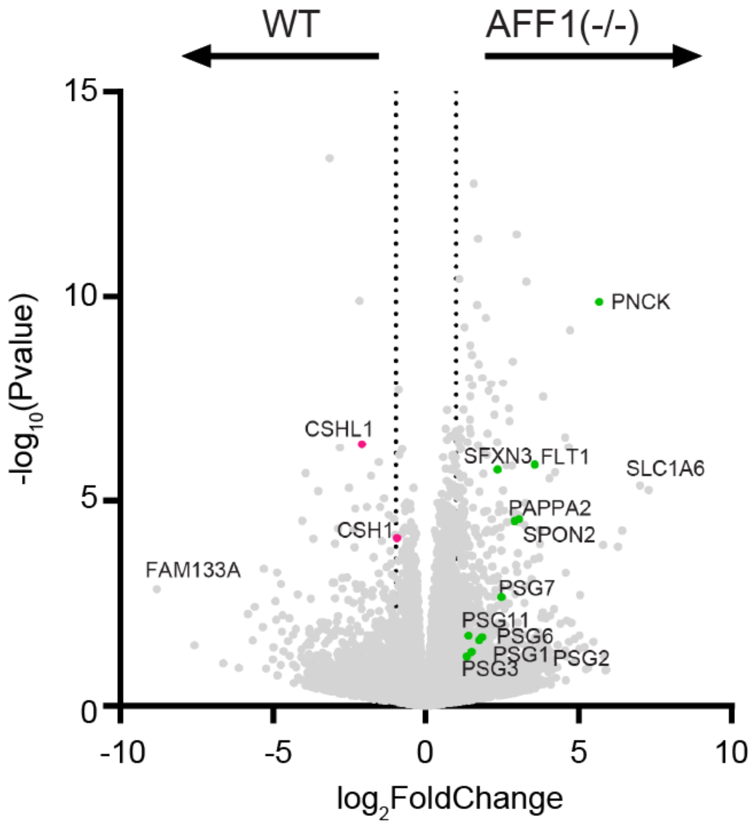


**Supplemental Figure 5.3. The deletion of AFF1 in TOs results in the downregulation of STB-2 marker genes.** Bulk RNA sequencing of WT or AFF1(-/-) $STB^{in}$ TOs was performed and analyzed with DESeq2 to determine differential gene expression. Dotted line represents a $log_2$ fold change greater than 1.

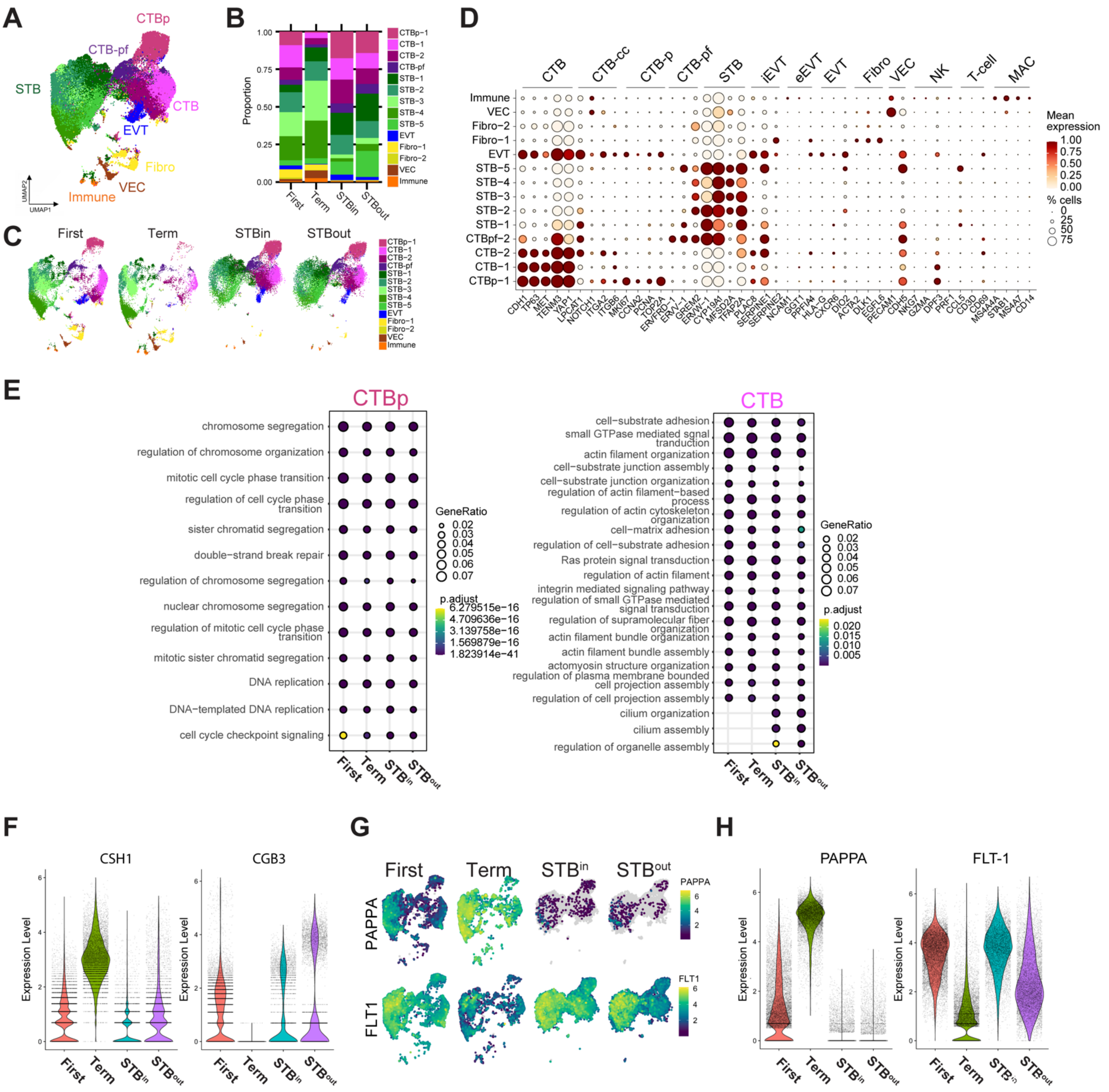

**Supplemental Figure 6.1: Characterization of nucleus types in merged SN dataset with TOs, first trimester, and full-term tissue. A** UMAP of the integrated SN dataset including first trimester, term tissue, STB$^{in}$ TOs, and STB$^{out}$ TOs. **B** Relative proportion of each nucleus type for each sample in the integrated dataset in A. **C** UMAP of each sample type split from the integrated dataset seen in A. **D** Dot plot of gene expression for the marker genes of each nucleus type of the entire integrated dataset, including nucleus types removed in A-C. The size of the dot demonstrates the percent of nuclei expressing a given gene and color represents the mean expression value. The labels on the top of the graph refer to the established marker genes used to identify the identity of each cluster. Nucleus types are annotated as follows: cytotrophoblasts (CTB), cytotrophoblast pre-fusion (CTB-pf), syncytiotrophoblast (STB), extravillous trophoblast (EVT), dendritic stem cell (DSC), vascular endothelial cell (VEC), fibroblast (Fib), natural killer cell (NK), and macrophage (MAC). **E** The top DEG for either CTB-p or CTB of each sample type were analyzed for GO term enrichment with clusterProfiler and plotted as a dotplot. CTB-pf was not analyzed due to the low number of CTB-pf cells found in full-term tissue. **F** Violin plot demonstrating gene expression of CSH1 and CGB3 in each sample type of the integrated dataset. **G** Feature plots demonstrating RNA expression of PAPPA and FLT-1. **H** Violin plot demonstrating gene expression of PAPPA and FLT-1 in each sample type of the integrated dataset.

**Supplemental Table 1**. Metadata of Tissues and TOs lines used in each experiment.

| Experiment | # of donors or TOs lines | Donor or TOs code | Gestational age | Donor or TOs sex | TO passage number |
|---|---|---|---|---|---|
| scRNA-seq of tissue | 3 | AGPL2 | 39 | M | n/a |
| | | AGPL3 | 39 | F | n/a |
| | | AGPL4 | 37 | M | n/a |
| snRNA-seq of tissue | 3 | AGPL3 | 39 | F | n/a |
| | | AGPL4 | 37 | M | n/a |
| | | AGPL5 | 38 | M | n/a |
| IF staining of tissue | 3 (6 independent tests) | AGPL3 | 39 | F | n/a |
| | | AGPL4 | 37 | M | n/a |
| | | AGPL5 | 38 | M | n/a |
| scRNA-seq of $STB^{in}$ TOs | 3 | TO68 | 27 w | M | P12 |
| | | TO81 | 40 w | F | P3 |
| | | TO82 | 40 w | M | P3 |
| snRNA-seq of $STB^{in}$ TOs | 3 | TO74 | 41 w | M | P5 |
| | | TO81 | 40 w | F | P2 |
| | | TO82 | 40 w | M | P3 |
| snRNA-seq of $STB^{out}$ TOs | 3 | TO74 | 41 w | M | P5 |
| | | TO81 | 40 w | F | P2 |
| | | TO82 | 40 w | M | P3 |
| snRNA-seq of $EVT^{enriched}$ TOs | 3 | TO74 | 41 w | M | P5 |
| | | TO81 | 40 w | F | P2 |
| | | TO82 | 40 w | M | P3 |
| IF staining in TOs | 2 (5 independent tests) | TO74 | 41 w | M | P10 |
| | | TO90 | 40 w | unknown | P3 |

**Supplemental Table 2**: Gene markers used for cell/nucleus type identification.

| Cell/Nucleus type | Cell/Nucleus type abbriviation | Markers |
|---|---|---|
| proliferative cytotrophoblast | CTBp | MKI67, CCNA2, PCNA, TOP2A |
| cytotrophoblast | CTB | CDH1, TP63, MET, TENM3, YAP1 |
| cytotrophoblast cell column | CTB-CC | LPCAT1, NOTCH1, ITGA2, ITGB6 |
| cytotrophoblast pre-fusion | CTB-pf | ERVFRD-1, ERVV-1, GREM2 |
| syncytiotrophoblast | STB | ERVW-1, CRYP19A1, MSFSD2A, TFAP2A |
| interstitial extravillous trophoblast | iEVT | PLAC8, SERPINE1, SERPINE2 |
| endothelial extravillous trophoblast | eEVT | NCAM1, GGT1, PPFIA3 |
| extravillous trophoblast | EVT | HLA-G, CXCR6, DIO2 |
| vascular endothelial cell | VEC | PECAM1, CDH5 |
| dendritic stem cell | DSC | PRL, IGFBP1, APOA1, SERPINA3 |
| natural killer cell | NK | NKG7, NCAM1, GZMA, DPF3, NKG7, NCAM1, PRF1 |
| T-cells | Tcell | CCL5, CD3D, CD69 |
| marcrophage | MAC | MS4A4A, STAB1, MS4A7, CD14 |
| fibroblasts | Fibro | ACTA2, DLK1, EGFL6 |

**Supplemental Table 3**. Composition of term trophoblast organoid medium (tTOM).

| **Ingredient** | **Final Concentration** |
|---|---|
| 100 × N2 | 1× |
| 50 × B27 | 1× |
| 500 × Primocin | 100 µg/ml |
| 80 × NAC | 1.25 mM |
| 100 × L-glutamine | 2 mM |
| 10000 × A83-01 | 500 nM |
| 10000 × CHIR99021 | 1.5 µM |
| 2000 × recombinant hEGF | 50 ng/ml |
| 2000 × recombinant hR-spondin1 | 80 ng/ml |
| 2000 × recombinant hFGF2 | 100 ng/ml |
| 2000 × recombinant hHGF | 50 ng/ml |
| 100 × Nicotinamide(NTM) | 10 mM |
| 200 × Y-27632 | 5 µM |
| 2000 × PGE2 | 2.5 µM |
| FBS | 10% (vol/vol) |
| Advanced DMEM/F12 | 1× |

**Supplemental Table 4.** Composition of EVT differentiation medium (EVTM).

| **Ingredient** | **Final Concentration** |
|---|---|
| 100 × L-glutamine | 2 mM |
| β-mercaptoethenol | 0.1 mM |
| 200 × penicillin/streptomycin | 0.5% (vol/vol) |
| BSA | 225 μg/mL |
| 100 × ITS-X supplement | 1× |
| NRG1 (w/o in EVT m2) | 100 ng/mL |
| A83-01 | 7.5 μM |
| Y-27632 | 2.5 μM |
| Knockout Serum Replacement | 4% (vol/vol) |
| Advanced DMEM/F12 | 1× |

**Supplemental Table 5:** sgRNAs sequence used for CRISPR/Cas9 mediated gene editing

| Target Gene | sgRNA pair | Sequence (5' → 3') | Target Strand | Target Exon |
|---|---|---|---|---|
| *RYBP* | **#1** | TCAGCTGGTGGCACAACAAG | Template strand | Exon 2 |
| | | ACCAGCTGAGAATTGATCCG | Coding strand | Exon 2 |
| | **#2** | TCGGATCAATTCTCAGCTGG | Template strand | Exon 2 |
| | | TTGAACATTTCAGAAAACCT | Template strand | Exon 2 |
| *AFF1* | **#1** | CCTGCTTCGAATTAGAGAGA | Coding strand | Exon 2 |
| | | GAAATGCCTCTTTCTCTTGG | Template strand | Exon 2 |
| | **#2** | CCTTCTCTCTAATTCGAAGC | Template strand | Exon 2 |
| | | AAATGCCTCTTTCTCTTGGT | Template strand | Exon 2 |

**Supplemental Table 6:** PCR primers used for sequencing validation

| Primer name | Sequence (5'->3') | PCR annealing temperature (℃) |
|---|---|---|
| RYBP_PF9 | GCAGACTGTATGCTGTTTGCTT | 60 |
| RYBP_PR9 | TACTGCACAGGGATGTGTATGTT | |
| AFF1_PF5 | ATGGCCCAGAGTGAATGCTT | 60 |
| AFF1_PR5 | ACTCAATTCCCCTGTTGAGAAGA | |

Additional Supplemental Files:

**Supplemental Table 7:** Primary and Secondary FISH probe sequences

File name: SupplementalTable7.xsl

**Supplemental Table 8:** Differentially expressed genes in each STB subtype in the $STB^{in}$+$STB^{out}$ integrated dataset.

File name: Supplemental_Figure3B_DEG.xlsx

**Supplemental Table 9:** GO terms and representative genes from each STB subtypes in the $STB^{in}$+$STB^{out}$ integrated dataset.

File name: SupplementalTable_Figure3C.xlsx

**Supplemental Table 10:** DEseq analysis of STB subtypes between $STB^{in}$ and $STB^{out}$ in the integrated dataset

File name: SupplementalTable_Figure3H_final.xlsx

**Supplemental Table 11:** DEseq analysis of bulk sequencing from the WT and knock out TO lines

File name: SupplementalTable_Figure5E.xlsx

**Supplemental Table 12:** Differentially expressed genes in the STB of $STB^{in}$, $STB^{out}$, first trimester tissue, and term tissue in the integrated Figure 6 dataset.

File name: SupplementalTable_Figure6A.xlsx

**Supplemental Table 13:** GO terms and representative genes from the STB of $STB^{in}$, $STB^{out}$, first trimester tissue, and term tissue in the integrated Figure 6 dataset.

File name: SupplementalTable_Figure6E.xlsx

**Supplemental Table 14:** DEseq analysis of STB between $STB^{in}$, $STB^{out}$ , first trimester tissue, and term tissue in the integrated Figure 6 dataset.

File name: SupplementalTable_Figure6F.xlsx
